# Proteomic profiling identifies co-regulated expression of splicing factors as a characteristic feature of intravenous leiomyomatosis

**DOI:** 10.1101/2022.03.17.484830

**Authors:** Lukas Krasny, Chris P Wilding, Emma Perkins, Amani Arthur, Nafia Guljar, Andrew D Jenks, Cyril Fisher, Ian Judson, Khin Thway, Robin L Jones, Paul H Huang

## Abstract

Intravenous leiomyomatosis (IVLM) is a rare benign smooth muscle tumour that is characterised by intravenous growth in the uterine and pelvic veins. Previous DNA copy number and transcriptomic studies have shown that IVLM harbours unique genomic and transcriptomic alterations when compared to uterine leiomyoma (uLM), which may account for their distinct clinical behaviour. Here we undertake the first comparative proteomic analysis of IVLM and other smooth muscle tumours (comprising uLM, soft tissue leiomyoma and benign metastasising leiomyoma) utilising data-independent acquisition mass spectrometry. We show that, at the protein level, IVLM is defined by the unique co-regulated expression of splicing factors. In particular, IVLM is enriched in two clusters composed of co-regulated proteins from the hnRNP, LSm, SR and Sm classes of the spliceosome complex. One of these clusters (Cluster 3) is associated with key biological processes including nascent protein translocation and cell signalling by small GTPases. Taken together, our study provides evidence of co-regulated expression of splicing factors in IVLM compared to other smooth muscle tumours which suggests a possible role for alternative splicing in the pathogenesis of IVLM.

## Introduction

Intravenous leiomyomatosis (IVLM) is a rare histologically benign smooth muscle tumour which is characterised by intravenous growth in the uterine and pelvic veins [1,2]. In some instances, it can extend into the inferior vena cava and the right heart which in rare cases may cause death [3,4]. IVLM is usually present with concomitant uterine leiomyoma (uLM) and one theory is that it originates from a pre-existing uLM where it extends and invades into the vessel wall [4,5]. Given that there are some instances where IVLM arises in the absence of a uLM [2,6], an alternate theory is that this tumour originates from the smooth muscle cells of the vessel wall. In addition to IVLM, there are other rare smooth muscle tumours with unusual quasi-malignant clinical behaviour such as benign metastasising leiomyoma (BML) and disseminated peritoneal leiomyomatosis [7,8].

Previous studies have undertaken comparative analysis of the molecular features of IVLM versus uLM to gain a better understanding of its underlying biology as well as the relationship between the two entities [9-15]. Some of the system-wide comprehensive profiling studies that have been reported include array comparative genomic profiling (aCGH) and transcriptomic analysis [9,11,13,14]. Collectively, these focused and system-wide studies indicate that IVLM share some cytogenetic and protein expression features with uLM (e.g. translocations in (12;14) and HMGA2 protein expression) [11,12,14,15], while at the same time harbour genetic and transcriptomic alterations that are unique. These unique alterations include distinct *MED12* mutations and elevated *HOXA13* gene expression in IVLM [10,12,13]. Given its rarity, all of the published Omics-based IVLM molecular profiling studies, with the exception of a recent study by Ordulu et al.[11], have been limited to a small number of cases (typically <5).

To date no proteomic profiling analyses have been undertaken in IVLM. Proteins are the critical drivers of cellular communication in normal cells and dysregulation of protein function is causative of many diseases including cancer [16,17]. We hypothesized that, unlike genomic and transcriptomic analysis, proteomic profiling will provide a more direct readout of the biological pathways and protein complexes that may play a role in the pathogenesis of IVLM [18,19]. Here we undertake a comparative mass spectrometry-based proteomic analysis of IVLM and other smooth muscle tumours (uLM, soft tissue leiomyoma (stLM) and BML), and demonstrate that at the protein level, IVLM is characterised by the unique co-regulated expression of splicing factors that comprise the spliceosome.

## Materials and methods

### Patients and tumour specimens

Use of archival formalin fixed paraffin embedded (FFPE) tumour samples and linked anonymised patient data was approved by Institutional Review Board as part of the PROSPECTUS study, a Royal Marsden-sponsored non-interventional translational protocol (CCR 4371, REC 16/EE/0213). One of the IVLM cases in this series has previously been described in a case report [20]. FFPE tissue from surgically resected primary tumours and accompanying annotation of baseline clinico-pathological variables were identified and retrieved through retrospective review of departmental database and medical notes at the Royal Marsden NHS Foundation Trust. The histological diagnosis was confirmed in all cases by experienced soft tissue pathologists (KT, CF). For each tumour, a single FFPE tissue block containing representative viable tumour was selected through review of haematoxylin and eosin (H&E)-stained sections. Five 20µm sections were cut from each selected tumour block and, where indicated, macrodissected to enrich to >75% viable tumour content.

### Protein extraction and sample preparation

The samples were processed as previously described [18]. Briefly, 20µm tissue sections from each sample were deparaffinised in xylene, rehydrated by washes with decreasing ethanol gradient and then dried. Samples were homogenized in lysis buffer (0.1M Tris-HCl pH 8.8, 0.50% (w/v) sodium deoxycholate, 0.35% (w/v) sodium lauryl sulphate) at a ratio of 200ul/mg of dry tissue using a LabGen700 blender (ColeParmer) with 3x 30s pulses. Homogenates were sonicated on ice for 10 min and then incubated at 95°C for 1 h to reverse formalin crosslinks. Lysis was continued by shaking at 750rpm at 80°C for 2 h. The resulting homogenate was then centrifuged for 15min at 4°C at 15,000rpm, the supernatant was collected and protein concentration of the supernatant was measured by bicinchoninic acid (BCA) assay (Pierce). The extracted proteins were digested using the Filter-Aided Sample Preparation (FASP) protocol as previously described [21]. Briefly, each sample was placed into an Amicon-Ultra 4 (Merck) centrifugal filter unit and detergents were removed by several washes with 8M urea. The concentrated sample was then transferred to Amicon-Ultra 0.5 (Merck) filters, reduced with 10mM dithiothreitol (DTT) and alkylated with 55mM iodoacetamide (IAA). The sample was washed with 100mM ammonium bicarbonate (ABC) and digested by trypsin (Promega, trypsin to starting protein ratio 1:100 µg) overnight at 37°C. Peptides were desalted on C18 SepPak columns (Waters), dried in a SpeedVac concentrator and stored at -80°C.

### SWATH-MS data acquisition and processing

Quantitative proteomic profiling was performed by sequential window acquisition of all theoretical fragments mass spectrometry (SWATH-MS) which is also known as data-independent acquisition mass spectrometry. Dried, desalted peptides were resuspended in a buffer A (2% ACN/ 0.1% formic acid), spiked with iRT calibration mix (Biognosys AG) and analysed on an Agilent 1260 HPLC system (Agilent Technologies) coupled to a TripleTOF 5600+ mass spectrometer with NanoSource III (AB SCIEX). 1 μg of peptides for each sample was loaded onto a self-made trap column packed with a 10 μm ReprosilPur C18AQ beads (Dr. Maisch) and washed for 5 minutes by buffer A. Peptides were then separated on a 75 μm×15 cm long analytical column with an integrated manually pulled tip packed with Reprosil Pur C18AQ beads (3 μm, 120 Å particles, Dr. Maisch). A linear gradient of 2–40% of Buffer B (98% ACN, 0.1% formic acid) in 120 min and a flow rate of 250 nl/min was used. Each sample was acquired in 2 technical replicates. Acquisition parameters were as follows: 60 precursor isolation windows with a fixed size of 13 Da across the mass range of m/z 380–1100 with 1 Da overlap. MS/MS scans were acquired in the mass range of *m/z* 100-1500. Cycle time of 3.1 s was used resulting in average 8 datapoints per elution peak. SWATH-MS spectra were analysed using Spectronaut 15.2 (Biognosys AG) against a published human library [22]. FDR was restricted to 1% on both protein and PSM level. Peak area of 3 to 6 fragment ions was used for peptide quantification. The mean value of max 6 peptides was used to quantify proteins while 2 unique peptides was set as a minimum requirement for inclusion of a protein in the subsequent analysis.

### Data processing and statistical methods

The proteomics dataset was further processed using R, Perseus 1.5.6 [23,24] and GraphPad 8.2.1. Protein quantities were log2 transformed and quantile normalised at sample level using proBatch package [25] in R followed by protein median centering across the samples. The normalized dataset was then visualized by hierarchical clustering using ComplexHeatmap package in R [26]. Gene Set Enrichment Analysis (GSEA) was applied using GenePattern online tool [27] to identify gene sets obtained from the MSigDB (c5.gobp.v7.5) [28] that were significantly enriched in IVLM samples. Similarly, single sample GSEA (ssGSEA) was applied using GenePattern to score sample-specific enrichment of the Spliceosome gene set from the KEGG pathways database [29]. To identify spliceosome components, the list of all identified proteins in this study was cross-referenced with the annotated spliceosome protein interaction dataset published by Hegele et al. [30]. Mutual co-expression of the splicing factors was assessed by Pearson’s correlation coefficient that was calculated in Perseus for all possible combinations of the identified splicing factors. The resulting similarity matrix was analysed and visualised by ConsensusClusterPlus [31] and ComplexHeatmap packages in R respectively.

To study association of the splicing factors identified in clusters 1-3 with known biological pathways, Pearson’s correlation coefficients between splicing factors and all other proteins in the proteomic dataset (after removal of all proteins annotated in the Spliceosome Database [32]) were calculated in Perseus. The resulting similarity matrices were hierarchically clustered and visualized by ComplexHeatmap package in R, where rows of each matrix were split into 4 clusters using k-means partitioning, Euclidean distance and 1000 repetitions. Subsets of proteins from the clusters with the highest and lowest average correlation were then used for over-representation analysis using DAVID 6.8 Functional analysis online tool [33].

## Results

### Quantitative proteomic profiling of smooth muscle tumours

The cohort is comprised of FFPE tumour material from 14 patients treated at The Royal Marsden Hospital. These specimens were obtained from surgical resections of IVLM (n = 3), uLM (n = 3), stLM (n = 7) and BML (n=1). Tumour specimens were subjected to sample preparation and protein extraction as depicted in Figure 1. Digested peptides then underwent proteomic profiling with SWATH-MS in technical duplicates. This analysis resulted in the identification and quantification of 2,473 proteins (Table S1). Unsupervised clustering of the full dataset shows that the IVLM cases largely cluster together separate from the stLM and uLM cases (Figure 2A). Interestingly the only BML case in the cohort clusters most closely to the IVLM cases.

**Figure 1:**
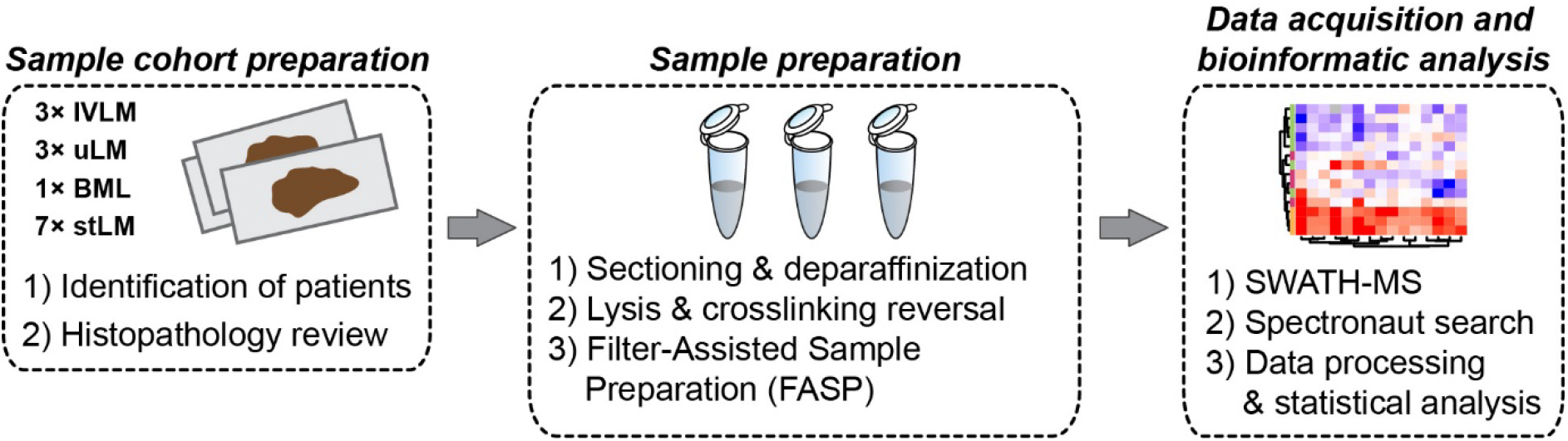
Experimental workflow depicting key procedures of sample selection and preparation, proteomic data acquisition and subsequent data processing and analysis.

**Figure 2:**
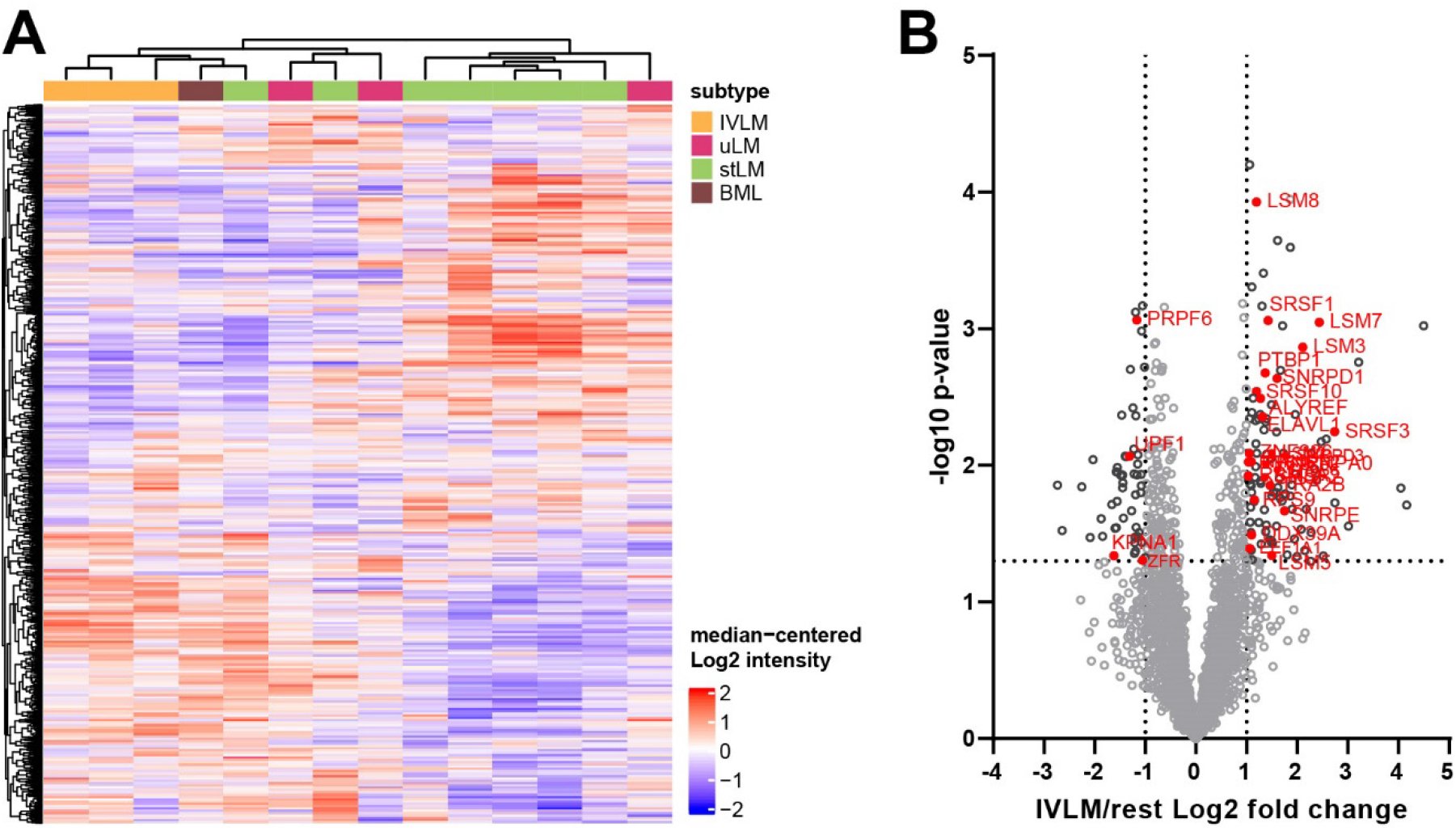
(A) Heatmap depicting unsupervised hierarchical clustering of 2,478 proteins that were quantified across all samples. The distance measure used for clustering is Pearson’s correlation. The full protein list is provided in Table S1. (B) Volcano plot depicting difference in protein expression between IVLM cases and all the other smooth muscle tumours (rest). Splicing factors with significantly different expression levels (>2 fold or < 2 fold) are highlighted in red.

Assessment of proteins that are significantly different in IVLM cases compared to uLM, stLM and BML cases identified 162 proteins of which 109 and 53 proteins are upregulated (>2 fold) or downregulated (<2 fold) in IVLM respectively (Fig 2B). Consistent with published immunohistochemical analysis studies [12], expression of the chromatin factor HMGA2, a protein which is highly expressed in IVLM due to the breakpoint on 12q14-15 [11,12,14], was not significantly different between IVLM and the other smooth muscle tumours in the cohort (Fig S1). Interestingly we find that 29/162 (18%) of the differentially expressed proteins are components of the spliceosome complex (Figure 2B).

### Enrichment of splicing processes in IVLM

To further investigate the biological processes that are enriched in IVLM compared to the other smooth muscle tumours, we undertook gene set enrichment analysis (GSEA) of the full proteomic dataset (Figure 3A). We show that the majority of the top 20 ranked enriched gene sets are processes associated with RNA splicing, processing, transport or metabolism. Beyond RNA-related biological processes, other enriched gene sets include protein targeting and localisation to membrane, regulation of gene transcription and translation. In line with the observation that a significant proportion of proteins enriched in IVLM are components of the spliceosome complex (Figure 2B), single sample GSEA (ssGSEA) of the proteomic data for each specimen in the cohort using the KEGG spliceosome gene set showed that the IVLM cases had significantly higher ssGSEA spliceosome scores compared to the other smooth muscle tumours in the cohort (Figure 3B). Taken together, our data indicate that both the spliceosome complex and biological processes involving RNA biology are enriched in IVLM specimens.

**Figure 3:**
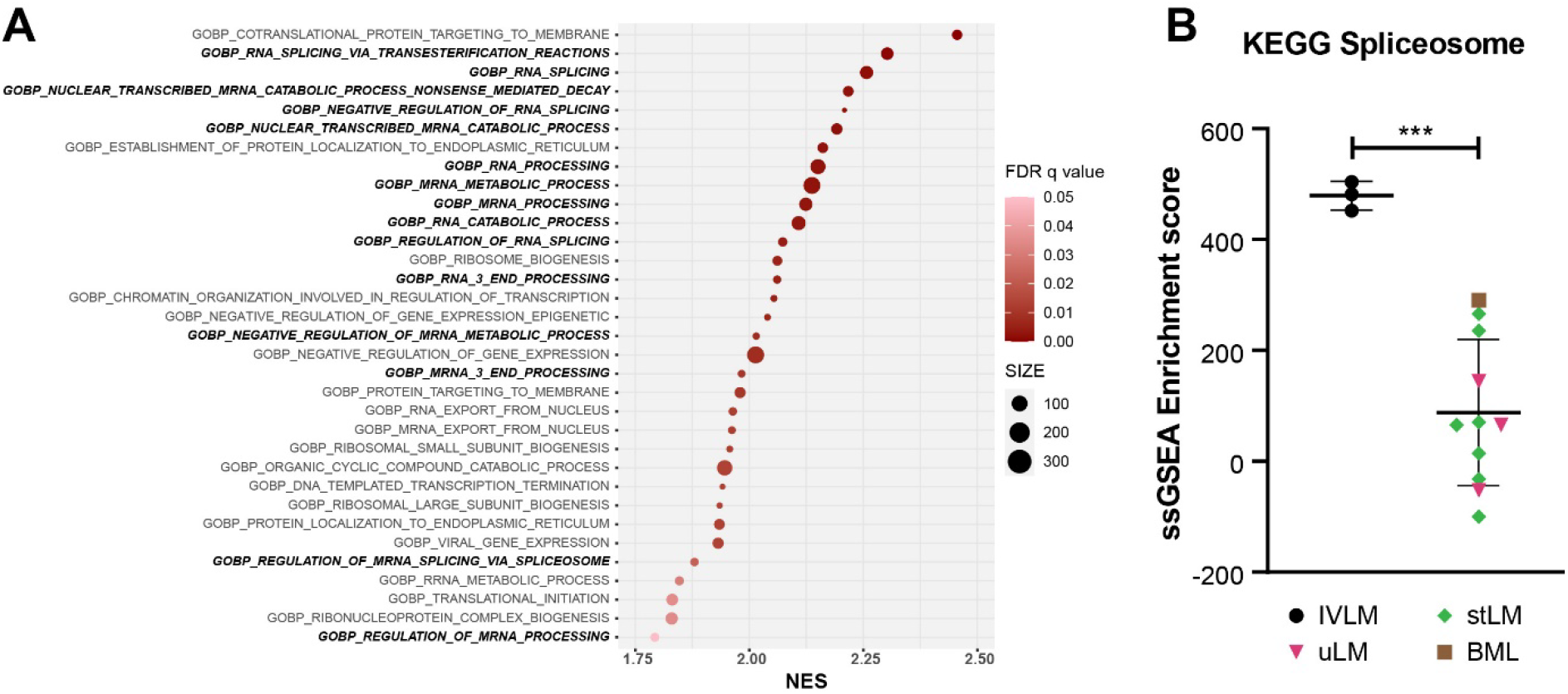
(A) Plot of Gene Set Enrichment Analysis (GSEA) results showing all the gene sets that are significantly enriched in IVLM samples. FDR q-value is represented by the colour of the circles while the size of the circles represents number of identified genes within each gene set. NES – normalized enrichment score. (B) Plot of single sample GSEA scores for the spliceosome gene set as defined by KEGG. The line and whiskers in plots represent mean and standard deviation. Statistical significance was calculated by two-sample t-test. *** p<0.001.

### Identification of co-regulated expression of splicing factors in the proteomic profiling dataset

It is well-established that the spliceosome is a highly dynamic macromolecular complex where more than 200 splicing factors are assembled into distinct complexes that vary in their composition in space and time [30,34]. We therefore hypothesized that despite the overall enrichment of spliceosome components in IVLM (Figure 3B), it is possible that subsets of co-regulated splicing factors may be responsible for the distinct clinical behaviour of IVLM versus leiomyomas. Indeed, unsupervised hierarchical clustering of 116 spliceosome components in the proteomic dataset showed that the spliceosome complex as a whole was not upregulated in IVLM (Figure 4A). Rather, there appeared to be subsets of splicing factors that were differentially expressed in IVLM, uLM and stLM.

**Figure 4:**
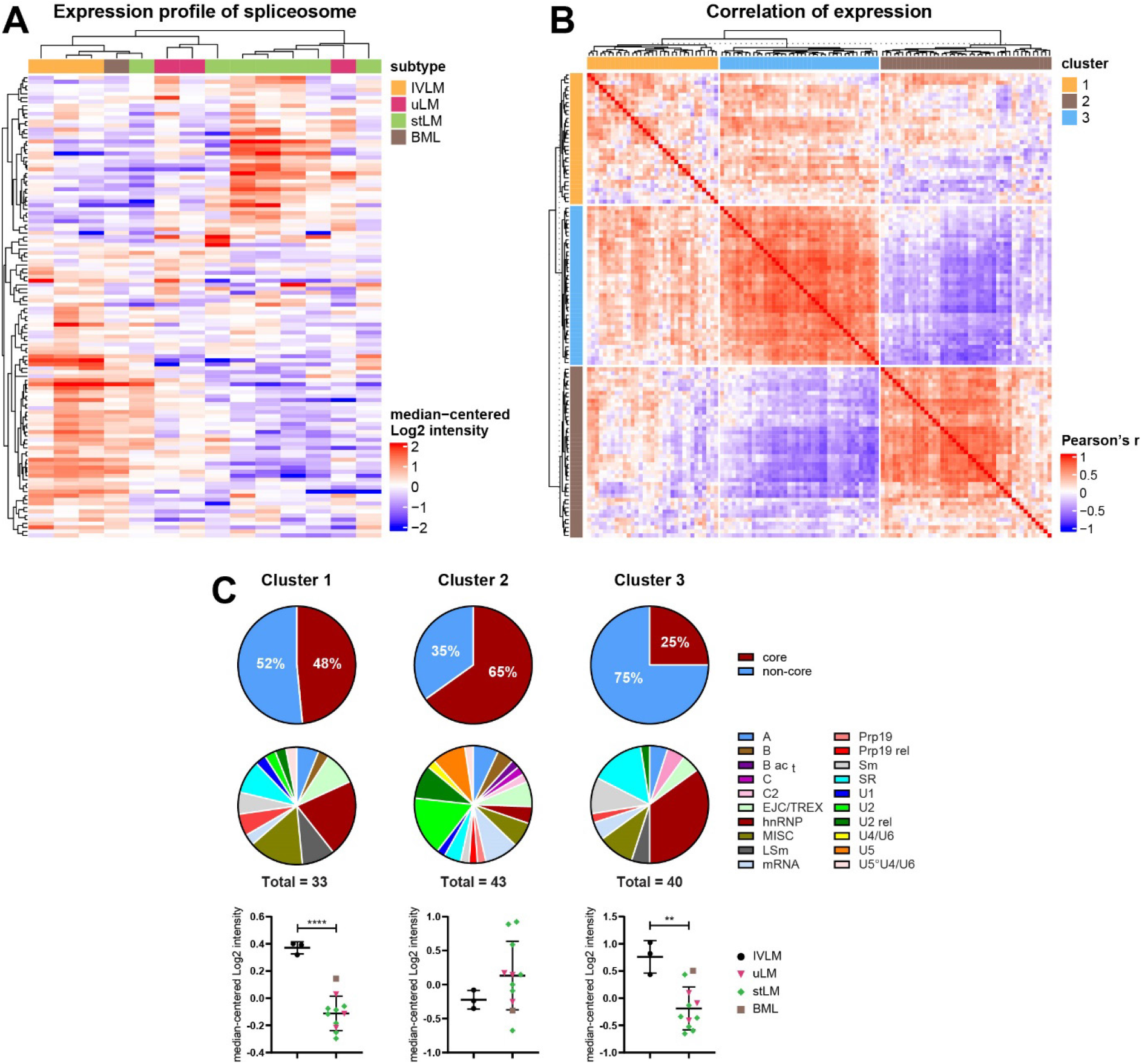
(A) Heatmap depicting unsupervised hierarchical clustering of 116 proteins of the spliceosome complex as defined by Hegele et al. [30]. The distance measure used for clustering is Pearson’s correlation. (B) Heatmap depicting similarity matrix of Pearson’s correlation coefficients of all possible pairwise combinations of the 116 splicing factors. Three clusters were identified by consensus clustering analysis. (C) Annotation and expression profile of the spliceosomal proteins belonging to clusters shown in Figure 4B. Venn diagrams depict spliceosome composition (core versus non-core, and distinct splicing factor classes) in each cluster while plots below show average expression levels of spliceosome components in each sample for a given cluster. Detailed composition of clusters and identity of individual proteins are listed in Table S2. The line and whiskers in plots represent mean and standard deviation. Statistical significance was calculated by two-sample t-test. ** p<0.01, **** p<0.0001.

Inspired by a previous study which showed that co-regulation of splicing factors is important in regulating breast cancer progression [35], we performed a Pearson’s correlation coefficient analysis of the protein expression levels of all possible combinations of 116 splicing factors in our dataset. Consensus clustering identified 3 clusters of splicing factors which is shown in the similarity matrix in Figure 4B (composition of each cluster provided in Table S2). In particular, Clusters 2 (n=43) and 3 (n=40) contain splicing factors which are negatively correlated between clusters but are positively correlated within clusters. Cluster 1 (n=33) is mixed with both positively and negatively correlated splicing factors.

### Distinct co-regulated clusters are comprised of splicing factors which are differentially expressed in IVLM versus the other smooth muscle tumours

An evaluation of the composition of splicing factors showed that each cluster is comprised of different proportions of core and non-core spliceosome proteins with Cluster 2 having the highest proportion of core proteins (65%) and Cluster 3 having the least core proteins (25%) (Figure 4C). Furthermore, assessment of the splicing factor classes based on nomenclature defined by Hegele et al., [30] finds that the splicing factor class composition of Clusters 1 and 3 is similar with the majority of proteins coming from the hnRNP, LSm, SR and Sm protein classes (Figure 4C). In contrast, the composition of cluster 2 is very different with U2, U2 rel and U5 protein classes dominating.

Quantitative assessment of the proteomic data showed that when broken down by cluster assignment, the IVLM specimens were significantly enriched in co-regulated splicing factors from Clusters 1 and 3 versus the other smooth muscle tumours in the cohort (Figure 4C). No significant difference between IVLM and the other smooth muscle tumours was seen in co-regulated splicing factors in Cluster 2. Collectively, this analysis indicates that at the protein level, IVLM is characterised by the co-regulated expression of specific classes of splicing factors that comprise the spliceosome.

### Co-regulated splicing factors are associated with multiple biological pathways, including protein translocation and signal transduction by small GTPases

We sought to determine if the expression of splicing factors in each of these clusters was linked to specific biological process. To do this, the Pearson’s correlation coefficient was calculated between all the proteins in the dataset (excluding spliceosomal proteins) and splicing factors in each of the three clusters. Unsupervised hierarchical clustering finds that 537 and 585 proteins were positively or negatively correlated with the splicing factors in Cluster 2, respectively (Figure 5A, clusters C and A). The same analysis in Cluster 3 identified positive and negative correlation in 545 and 738 proteins, respectively (Figure 5B, clusters C and B). Unsurprisingly, since Cluster 1 comprised of both positively and negatively correlated splicing factors, no significantly correlated proteins were found in our dataset (data not shown). Given that Clusters 2 and 3 have opposing profiles in co-regulated splicing factors (Figure 4B), it is expected that proteins correlating with these clusters would follow the same trend. Indeed, we demonstrate that there was substantial overlap of proteins which show opposite co-expression patterns (i.e. positively correlated proteins in Cluster 2 and negatively correlated proteins in Cluster 3), and vice versa (Figure 5C).

**Figure 5:**
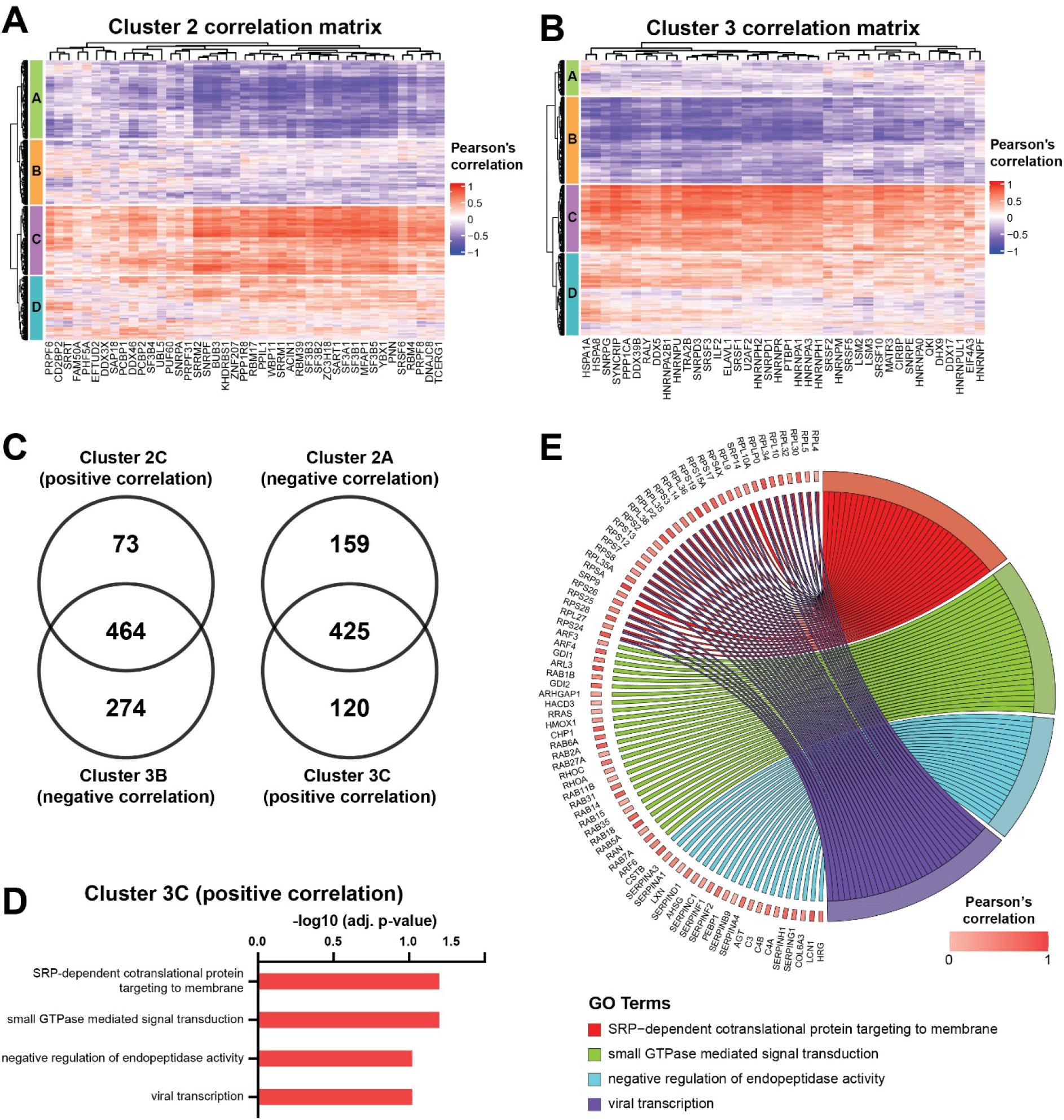
Heatmaps depicting correlation matrix of Pearson’s correlation coefficient calculated between the splicing factors in (A) Cluster 2 or (B) Cluster 3 and all the other proteins in the dataset that are not part of the spliceosome complex. Heatmaps are split into four clusters based on k-means partitioning. (C) Venn diagrams depicting the overlap between the positively and negatively correlated proteins in Cluster 2 and 3 respectively, and vice versa. (D) Plot of overrepresentation analysis results showing ontologies which are positively correlated with the splicing factors in Cluster 3 (FDR < 0.1). (E) Chord plot depicting all positively correlated proteins identified by overrepresentation analysis in Fig 5D.

Focusing on Cluster 3 which is significantly upregulated in IVLM (Figure 4C), over-representation analysis finds 4 ontologies that are enriched in the proteins that are positively correlated with the splicing factors in this cluster (Figure 5D). These ontologies include nascent protein targeting to the endoplasmic reticulum (SRP-dependent cotranslational protein targeting to membrane), signal transduction mediated by small GTPases, hydrolysis of proteins by peptidases (negative regulation of endopeptidase activity) and proteins involved in viral transcription. The positively coregulated proteins in these ontologies is shown in the chord diagram in Figure 5E.

## Discussion

IVLM is a rare benign smooth muscle tumour with quasi-malignant clinical behaviour. Previous profiling studies characterising its molecular features have focused on DNA copy number and transcriptomic alterations [9,11,13,14]. Here we performed the first proteome level analysis of IVLM and compare it to other smooth muscle tumours including uLM, stLM and BML. We show that IVLM is characterised by a differential expression of spliceosome complex components. In particular, by utilising a bioinformatics approach to delineate co-regulation of splicing factors, we find that there are two specific clusters of co-regulated splicing factors in the hnRNP, LSm, SR and Sm protein classes that are enriched in IVLM compared to the other smooth muscle tumours in this cohort. Finally, we demonstrate that one of these clusters (Cluster 3) is associated with high expression of proteins involved in key biological processes such as nascent protein translocation and signalling by small GTPases. To our knowledge this is the first demonstration that IVLM is characterised by a distinct group of co-regulated splicing factors, which may contribute to its unique clinical behaviour. It highlights the utility of proteomics to provide novel insights into IVLM tumour biology beyond the current state-of-the-art gained from published aCGH and gene expression studies.

Splicing occurs through a complex series of well-regulated steps mediated by the spliceosome machinery [36]. It has been shown that aberrations in specific splicing factors disrupt the composition of the spliceosome complex and drive carcinogenesis [37,38]. For instance, mutations in the splicing factor SF3B1 in both solid and liquid cancers initiate oncogenic alternative splicing reprogramming that is key to cancer development and progression [39-43]. Furthermore, it has been recently shown that some of these splicing factor mutations may induce new vulnerabilities that can be therapeutically exploited in a synthetic lethal fashion [44-46]. In the same vein, it is possible that the distinct co-regulation of splicing factors observed in IVLM may result in dysregulated alternative splicing that could account for its intravenous growth patterns. Unfortunately, due to the highly fragmented nature of total RNA extracted from FFPE specimens, we were unsuccessful in our efforts to measure alternative splicing profiles by RT-PCR from the cases in this series despite multiple repeated attempts. Future RNASeq or RT-PCR analysis on prospectively collected flash frozen specimens would be key to establishing if differential alternative splicing occurs in IVLM versus uLM. Identifying such alternatively spliced genes could offer a mechanistic explanation into the quasi-malignant behaviour of IVLM.

This study is limited by the small number of IVLM cases that were studied. IVLM is a rare condition and the vast majority of profiling studies to date comprise a small number of cases (typically <5). Despite the limited numbers, we were able to demonstrate that there was a statistically significant enrichment of co-regulated spliceosome components in IVLM. Interestingly, we show that the sole BML case in our cohort clustered most closely to the IVLM cases (Figure 2A). BML is another rare unusual variant of leiomyoma that often manifests as multiple nodules in the lungs and other sites [47]. A recent aCGH analysis finds that IVLM and BML share recurrent copy number alterations that are rarely seen in uLM [11]. Consistent with this finding, our data shows that at the proteomic level, BML is more similar to IVLM compared to uLM. It is however important to note that our proteomic analysis was performed on a small case series treated within a single institution and any findings will need to be independently validated.

## Conclusions

In summary, we have undertaken a comparative proteomic profiling study of IVLM and other smooth muscle tumours (uLM, stLM and BML) and describe the selective enrichment of co-regulated splicing factors which are associated with distinct biological pathways. We anticipate that future work integrating proteomics with complementary Omics-based profiling approaches such as RNAseq will shed further insights into the possible role of alternative splicing in the pathogenesis of IVLM.

## Institutional Review Board Statement

The study was conducted in accordance with the Declaration of Helsinki, and approved by the Institutional Review Board of The Royal Marsden NHS Foundation Trust (PROSPECTUS study, CCR 4371, REC 16/EE/0213)

## Informed Consent Statement

Informed consent was obtained from all subjects involved in the study.

## Data Availability Statement

The mass spectrometry proteomics data have been deposited to the ProteomeXchange Consortium via the PRIDE [48] partner repository with the dataset identifier PXD031637.

## Supplemental figure

**Figure S1:**
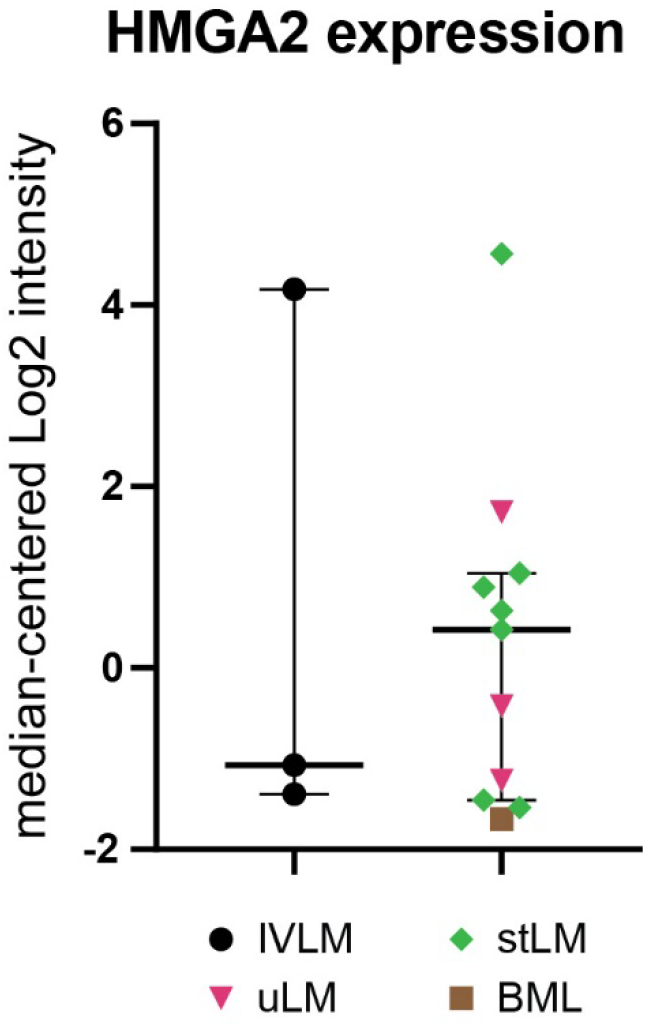
Expression levels of HGMA2 protein of each case in the cohort. The line and whiskers in plots represent mean and standard deviation. Expression levels of this protein was not significantly different between IVLM and the other smooth muscle tumours in the cohort

## Supplemental table legends

**Table S1:** Full proteomic dataset for the cohort. The dataset was log2 transformed and quantile normalized. Reported values represent median centred protein expression levels.

**Table S2:** List of splicing factors found in the three individual consensus clusters which are annotated based on spliceosome complex protein classes as defined by Hegele *et al*., Mol. Cell. 2012

